# Bow-tie motifs enable protein multifunctionality by connecting cellular processes in the interactome

**DOI:** 10.1101/495895

**Authors:** Kristoffer Niss, Jessica X. Hu, Cristina Gomez-Casado, Thorsten Joeris, William W. Agace, Kirstine G. Belling, Søren Brunak

**Affiliations:** Novo Nordisk Foundation Center for Protein Research, Faculty of Health and Medical Sciences, University of Copenhagen, DK-2200 Copenhagen, Denmark; Section of Biology and Chemistry, Department for Micro- and Nanotechnology, Technical University of Denmark, DK-2800 Kgs. Lyngby, Denmark; Mucosal Immunology, Faculty of Medicine, Lund University, SE 221 00 Lund, Sweden; Institute of Applied Molecular Medicine (IMMA), Faculty of Medicine, San Pablo CEU University, 28925, Madrid, Spain

**Keywords:** Network motifs, bow-tie, cell types, protein interactome, functional organization

## Abstract

Cell type-specific protein interactomes are promising resources for the understanding of cell functionality and genetic perturbations in cellular states. However, while many low-and top-level aspects of interactome topology have been mapped, we still lack an understanding of the topological motifs at the intermediate level, where connectivity between cellular processes takes place. We combine conventional dendritic cell lineage 1 (cDC1)-specific expression data with a high-quality protein interactome, subsequently creating an information-rich cDC1-specifc interactome matrix that displays topological information on an unprecedented 36.7 million protein pairs at once. By dissecting the interactome-matrix, we reveal that the bow-tie motif is a major connector of cellular processes and an extremely widespread topology in the protein interactome. In aggregation, the bow-tie motifs form a scaffold within the interactome that hierarchically link cellular processes together. We further demonstrate that the bow-tie motif’s design can enable protein multifunction. Our generic approach allows a topological characterization and visualization of any large network, facilitating a more effective usage of community-produced interaction data, and is applicable to any cell type’s interactome.

## Introduction (638 words)

Distinct cellular states, such as apoptosis, differentiation or migration, drive cellular processes. These states cannot be adequately described by the activity of a single cellular process, like sugar metabolism or antigen processing, but only by an ensemble of cooperating cellular processes. The notion of network biology – cooperation and connectivity between cellular processes – is generally accepted, exemplified by singular perturbations that can disrupt multiple cellular processes in concert through propagation via protein, RNA or metabolite interactions [1]. Just as a protein working in isolation has limited impact on system functionality, in cellular processes a similar principle applies. Thus, to understand a cellular state and how it may be dysfunctional, both the implicated cellular processes underlying that state, and their inter-connectivity, must be considered.

Comprehensive interactomes of high-quality protein-protein interactions (PPIs) have proven valuable for studying the inter-connectivity of cellular processes [2–4]. In such studies, cellular processes are linked to their respective molecular machinery, i.e. protein complexes, increasingly often in a cell type-specific interactome context [5,6]. Interactions and patterns of perturbation propagation can thereby be identified [7–9]. This has uncovered multiple connections between cellular processes [10] and identified sets of proteins facilitating these connections. These are known as, among others, interface-and extreme multifunctional proteins [1,10,11]. However, it has turned out to be difficult to construct a comprehensive overview of which cellular processes are definitively connected with which, and when.

A major challenge is the insufficient understanding of the organization of the protein interactome, i.e. topology, at the level of cellular processes. The topology of the protein interactome can be divided into at least three organizational levels: top, intermediate and low (**Figure 1**). At the top level, considering the whole interactome, studies have revealed a hierarchical structure containing modular organizations of proteins [12,13]. At the low level, considering structures of ∼3-10 proteins, small topological motifs such as bi-fans or loops constitute the topological organization [14]. However, at the intermediate level, where cellular processes and their inter-connectivity reside, the topology is largely unknown.

Mapping the topology at the intermediate level is essential in order to learn which cellular processes are connected, and how, and when. This will improve our general understanding of protein interactomes, and facilitate the unraveling of how network perturbations disrupt multiple cellular processes in concert.

**Figure 1:**
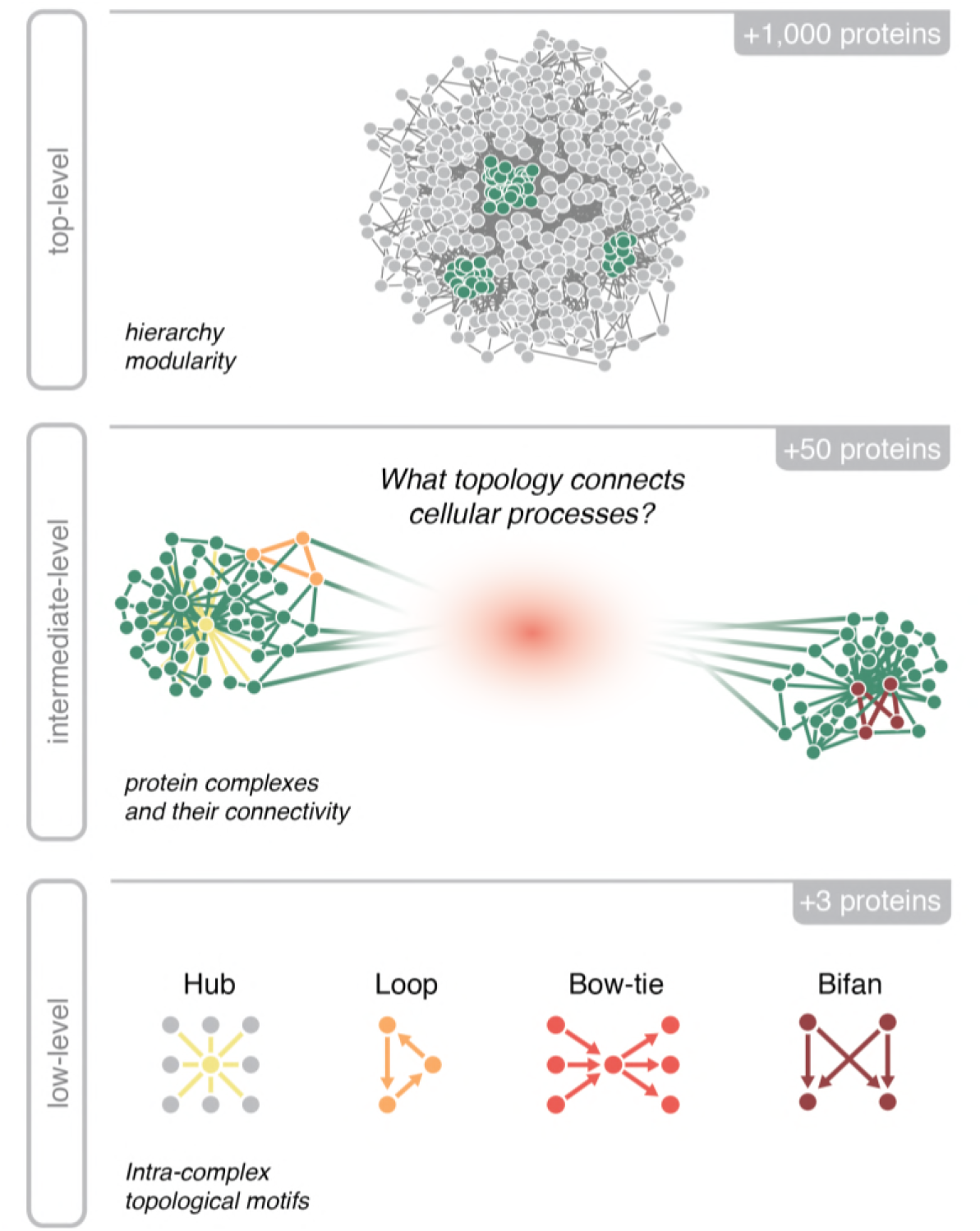
A protein interactome’s organizational levels. The topology of a protein interactome can be divided into at least three levels: top, intermediate and low. The top level describes the protein interactome as a whole, typically as a hierarchical structure that connects modular organizations of proteins, i.e. protein complexes. The intermediate level is the description of the topology between protein complexes and their exact inter-connectivity layout. This level is poorly characterized. The low level constitutes small topological motifs such as hubs or loops. These structures are often found within the larger protein complexes.

The importance of topological motifs is further emphasized by their inherent features, such as robustness or regulatory power, which optimizes them for different functional roles [15–17]. Consequently, when a certain topological motif is found at a specific location in the protein interactome, it will hint at the functional tasks of that location. This task could be modifiable signaling, feedback regulation or something else. The hub protein is an example of a motif with a specific function, as it is often located where extensive regulation is needed [7].

We present a general approach for untangling the topology of any large network, and apply it to the intermediate topological level of the conventional dendritic cell lineage 1 (cDC1)-specific interactome. We visualize the topology of 36.7 million unique protein pairs at once in a cDC1-specifc interactome matrix. Based on the matrix, we identify topological motifs, such as protein complexes and topologies that are situated between them. These large topological motifs enable us to systematize the 278,365 cDC1-specific PPIs simultaneously and in effect filter away PPIs that deviate from prevalent topological trends. We find the bow-tie motif [15,16] to be a significant connector of cellular processes, and analyze its functional role at the intermediate level. The bow-tie motif is abundant, hierarchically connects cellular processes and displays repetitive connections between specific processes. Furthermore, we find that the proteins centrally located in the bow-tie, i.e. the “knot” proteins, are multifunctional. The multifunctionality of “knot” proteins is probably assisted by the bow-tie motif’s design, which supports easy recruitment of the “knot” proteins to multiple distinct cellular processes.

## Results

### Construction and visualization of the cDC1 interactome

To study the intermediate level of a protein-protein interactome in a cell type-specific context, we constructed an interactome specific to cDC1. All actively expressed genes in cDC1 cells (see definition in Methods) were identified by generating and analyzing RNA-seq data obtained from FACS sorted XCR1^+^MHCII^hi^ cDC1 extracted from murine mesenteric lymph nodes (**Supplementary Figure 1**). This data-type had deep transcriptomic coverage, while it retained cell specificity. We built the cDC1-specific interactome by sub-setting the physical PPI database InWeb_IM [2], keeping only proteins with genes expressed in cDC1 cells (n=8,569). InWeb_IM is an aggregation of experimentally generated PPI datasets and databases from human, mice and other eukaryotes. Each InWeb_IM interaction has been quality benchmarked and scored accordingly [2].

To enable a topological “autopsy”, we visualized the cDC1-specific interactome in its entirety (**Figure 2A**). We portrayed the full interactome using a highly information-rich matrix that displays the topological information on every one of the 36.7 million unique protein pairs (73.4 in the symmetric matrix), corresponding to a 73-megapixel image. This novel and comprehensive figure shows the direct high-confidence PPIs in the upper matrix and the so-called weighted topological overlap (wTO) in the lower matrix [13,18]. The wTO is, in simple terms, the relative number of shared interactors between two proteins (see **Figure 2B** and Methods for details). We used the wTO measure to hierarchically cluster the matrix since the wTO captures more topological information than the scored direct interaction alone. **Figure 2A** is of interest in itself, as it is, to our knowledge, the first comprehendible visualization of the entire topology of a cell type-specific protein interactome.

**Figure 2:**
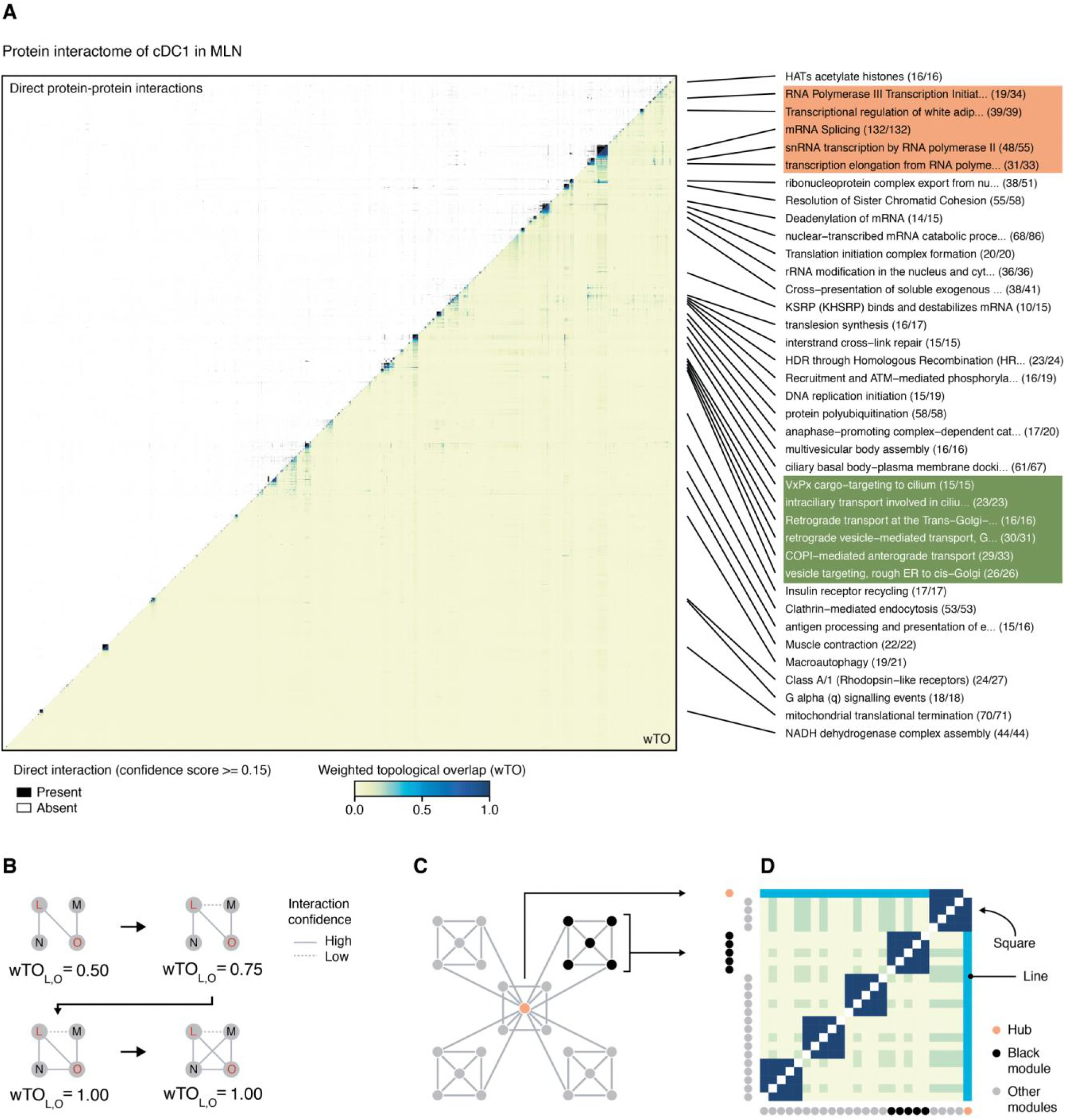
Network topology of the cDC1-specific protein interactome. (A) In the upper matrix, direct protein-protein interactions from InWeb_IM are displayed in black. In the lower matrix, the weighted topological overlap (wTO) measure (explained in (B)) between protein pairs is displayed. To the right, the larger protein complexes (n ≥ 15) are annotated using the Gene Ontology Biological Process database or Reactome. In parentheses, we indicated the number of proteins significant for the annotation term relative to the complex size. The matrix is hierarchically clustered based on the wTO measure. Squares in orange (transcription) and green (intracellular transport) highlight areas where related cellular processes clustered together. **(B)** Graphical visualization of the wTO measure. The wTO measure increases as proteins become more inter-connected, here exemplified by protein L and O: wTO_L,O_ increases to 0.75 when L-M connects and further increases to 1.0 when also N-O connects. Yet, wTO_L,O_ does not increase by adding the N-M connection since this connection does not increase the inter-connection between L and O. **(C and D)** A mock network and its corresponding wTO matrix. The mock network contains two types of topological motifs: A hub (orange node) and five modular structures (grey/black nodes). Correspondingly, in the wTO matrix, the orange hub motif is observable as long horizontal and vertical lines, while the modular motifs appear as dark blue squares. Thus, the interactome topology can be understood from the matrix by linking specific wTO features to known topological motifs.

### Identifying protein complexes using the wTO matrix

The cDC1 interactome matrix in **Figure 2A** encodes the interactome topology at all levels. The topology can be understood by decoding the substructure of the matrix. To demonstrate the decoding process, we generated a mock network and its corresponding wTO matrix (**Figure 2C-D**). Using this, we show that the well-known topological motifs, hub proteins and modular protein organizations, create specific matrix features of long lines and dark squares, respectively. By coupling recognizable wTO features to specific topological motifs we can backtrack, and hereby learn the topology of even very large networks directly from the wTO matrix.

The intermediate level of the cDC1 interactome consists of protein complexes and their topological connections. To describe the intermediate level, we first defined all protein complexes seen as squares in the cDC1 interactome matrix. By computationally traversing the cDC1-specific matrix of direct PPIs along the diagonal, we tagged and counted all protein complexes (**Supplementary Figure 2A**). In total, we tagged 64 protein complexes in the matrix, where each protein complex contained between 9-132 protein members.

We made sure that each protein complex was significantly linked to a specific cellular process by annotating each complex with its most significant term from either the *Gene Ontology (GO) Biological Process Database* or the *Reactome Database*. The 64 protein complexes were all significantly enriched for multiple annotations and the most significant annotation was, for most of the complexes, annotated with very high precision (**Supplementary Figure 2B**). Similarly, we inspected how the protein complexes were arranged in relation to each other. Previous studies have shown that protein complexes with related cellular processes are close in proximity in the protein interactome [12,15]. We also found this to be the case in the cDC1 interactome matrix, illustrated by two separate clusters of protein complexes. Each cluster contained multiple cellular processes that were all related to the same high-level process (**Figure 2A**, *transcription cluster* colored orange and *intracellular transport cluster* colored green).

### Topological motifs between protein complexes

Having identified the protein complexes at the intermediate level, we turned to the topological motifs that connected them. In the cDC1 interactome matrix, the topology between protein complexes is any matrix feature that is situated between them. Though we mostly observed a scatter of interactions in that area, we found one recurrent feature: vertical and horizontal *dashes*. We present examples of these dashes colored red in **Figure 3A** (subdomain of **Figure 2A**). The dashes were distinct from the line feature of hub motifs, as they were darker colored and shorter in length. To investigate the dash feature, we extracted two protein complexes from **Figure 3A** (in blue), which had 11 dashes between them, and presented them as a graph in **Figure 3B**. From this example, we saw that the dash features were the topological motif, the bow-tie [15–17,19,20]. For the remainder of this work, we define the simplest bow-tie motif as a single “knot” protein suspended between two fans of interactions (**Figure 3B, Example**).

**Figure 3:**
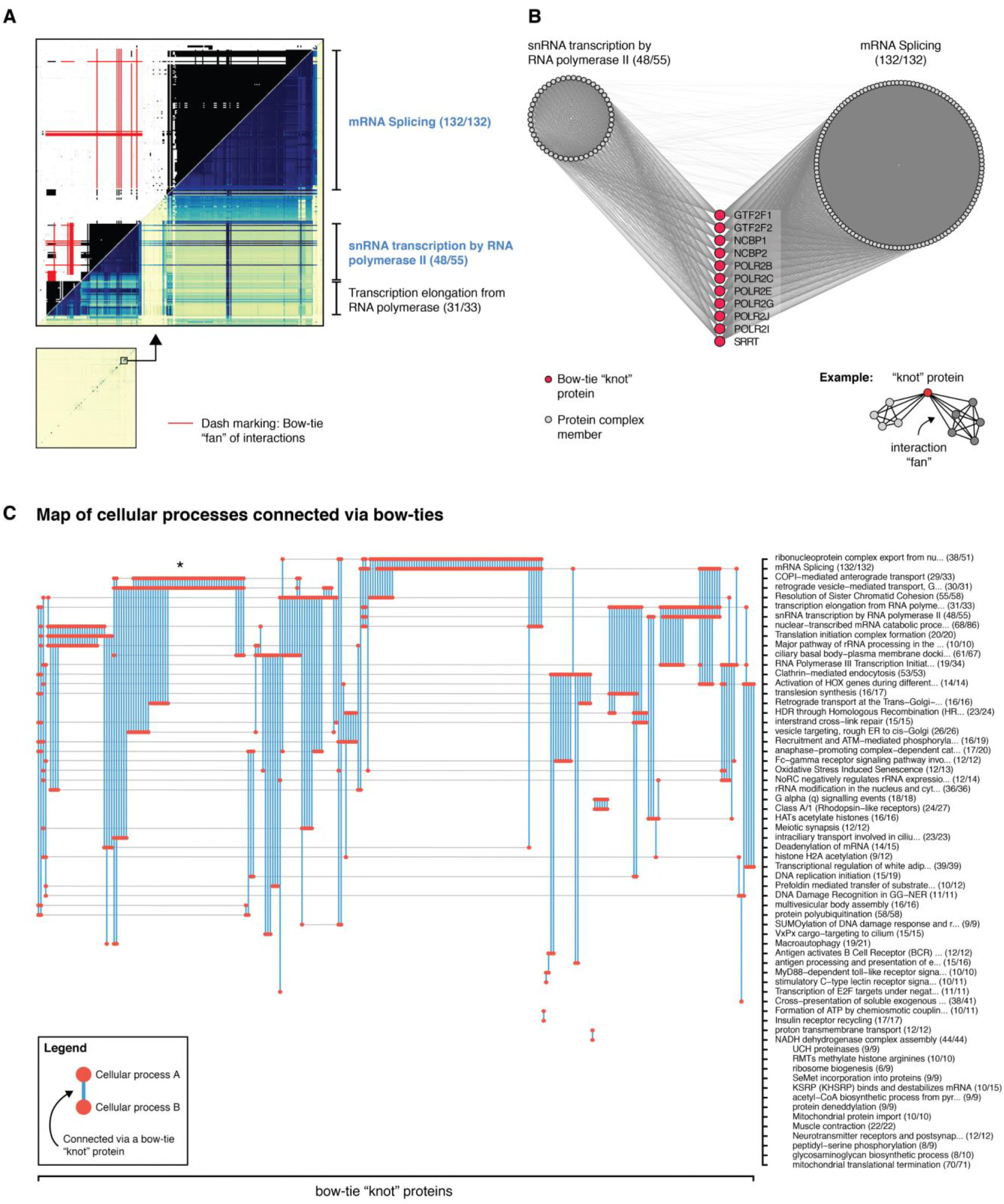
Bow-tie “knot” proteins connect fundamental cellular processes. (A) Subdomain of the wTO matrix in Figure 2A to highlight the dash features, here colored red in the upper matrix. **(B)** The two protein complexes named in blue in (A) visualized as a network. The eleven dashes in (A) are here identified as interaction fans of eleven bow-tie motifs suspended between the two protein complexes. **(C)** A map of cellular processes linked by bow-tie motifs. The 294 identified “knot” proteins are hierarchically ordered along the x-axis and annotated protein complexes are similarly ordered along the y-axis. The vertical blue lines with red dots show protein complexes connected via bow-ties. Note, many “knot” proteins connect more than two protein complexes, so more than two dots can occur on a blue line. Indented protein complexes low on the y-axis are not connected by bow-ties. We find that especially fundamental cellular processes utilize the bow-tie connections and that some pairs of cellular processes have repeated connections (**asterisk**).

### Bow-tie motifs connect cellular processes hierarchically and repeatedly

To obtain an overview of the frequency of the bow-tie motif in the cDC1 interactome, we catalogued the complete set of bow-ties and “knot” proteins. This was done computationally by searching the direct interaction matrix of cDC1 (**Figure 2A**), tagging all dash features created by bow-tie motifs, resulting in a final count of 294 “knot” proteins (**Supplementary Figure 3**). We hereafter constructed a map of the bow-tie facilitated connectivity between cellular processes (**Figure 3C**). In total, 51/64 cellular processes were connected by bow-ties. At closer inspection, we found that bow-tie motifs connected specific cellular processes in a repetitive manner, where multiple bow-ties were suspended between the same two cellular processes (**Figure 3C, asterisk**). Furthermore, the organization of the bow-tie links appeared hierarchical. For example, the complexes “COPI-mediated anterograde transport” and “Retrograde vesicle-mediated transport” were repeatedly connected, thus forming a “core” (**Figure 3C, asterisk**). Other complexes, such as “Retrograde transport at the Trans-Golgi-Network” and “Vesicle targeting rough ER to cis-Golgi”, were also connected to the core, but less frequently, hereby forming a hierarchy.

The map also revealed key aspects of the functional organization of cellular processes. We found that especially fundamental cellular processes like DNA repair, transcription, splicing or intracellular vesicle transport were often repeatedly connected to each other by bow-ties. Conversely, cellular processes linked by few or no bow-ties were, in general, either immune cell related (e.g. *antigen activates B Cell Receptor (BCR) leading to generation of second messengers*), independent molecular entities (e.g. *mitochondrial translational termination*), or smaller protein complexes of less than ten proteins.

The many bow-tie connections between fundamental cellular processes suggested to us that the bow-tie motifs were not specific to cDC1, but perhaps a component of all interactomes, independent of cell type. Thus, we checked the expression of the 294 cDC1 “knot” proteins in six other types of immune cells in a separate public dataset, sorted from blood from healthy human donors [21]. We found that 81-97% (Median: 93.5%) of the “knot” proteins were expressed in all six cell types (**Supplementary Table 1**), which further indicated that the bow-tie motif may be a general topological component of cellular protein interactomes.

### Functional context of the bow-tie motif

The bow-tie motif is by design a powerful “signal coordinator” [16,17,19]. A signal may arrive from alternating protein receptors (bow-tie; in-fan). The central “knot” molecule, often a secondary messenger [15], then coordinates the input signal and translates it to a steady output (bow-tie; out-fan). Hereby, the bow-tie motif dampens input noise, which is necessary in fluctuating biological networks. At least one bow-tie in our catalogue is a known “signaling coordinator”, i.e. the TRAF6-IRAK bow-tie core in the “MyD88-dependent TLR signaling pathway” [20] (shown as a network in **Supplementary Figure 4**).

However, we still speculated as to whether all 294 “knot” proteins acted solely as “signal coordinators”. Looking into this, we found that at least a third of our “knot” proteins performed molecular functions not typically involved in signal transduction (**Figure 4A**). Furthermore, 100/294 “knot” proteins had more than two interaction fans (**Figure 4B**), which does not comply with the input/output format of the standard bow-tie “signal coordinator”.

**Figure 4:**
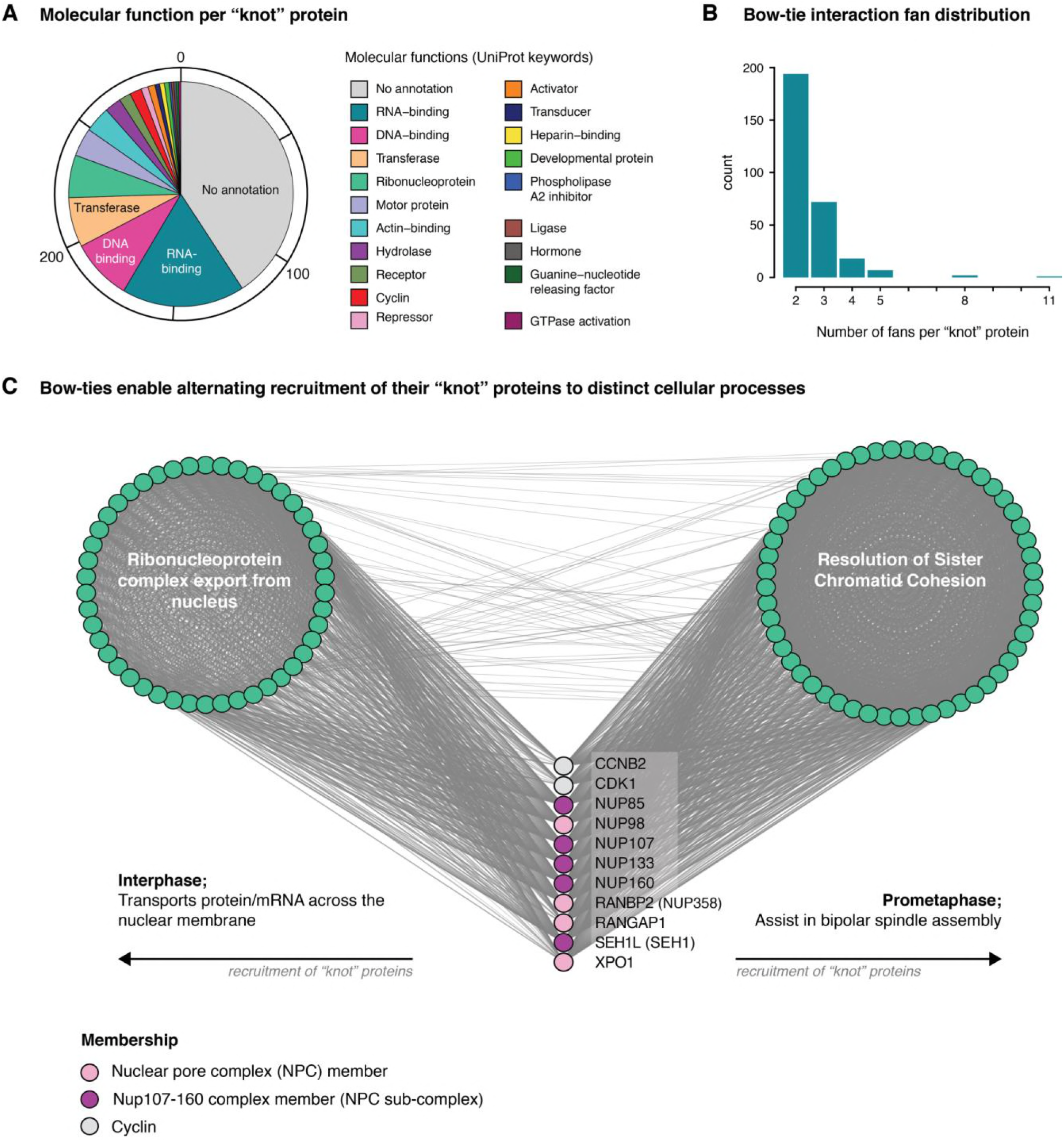
Functional aspects of the bow-tie motif. (A) Pie chart showing the main molecular function of each of the 294 “knot” proteins. It demonstrates that many “knot” proteins perform roles not typically linked to signaling cascades, such as RNA-binding, DNA-binding and acting as ribonucleoprotein subunits. **(B)** Distribution of the number of interaction fans per “knot” protein. In total, 100 “knot” proteins have more than two fans, which is atypical for bow-ties in signaling cascades. **(C)** Network representation of the cellular processes “Ribonucleoprotein Complex Export from Nucleus” and “Resolution of Sister Chromatid Cohesion” with eleven “knot” proteins suspended via bow-tie motifs between the them. The bow-tie motifs appear to enable the “knot” proteins (pink, purple and grey) to be recruited by distinct cellular processes depending on cell need. The bow-ties in (C) do not appear to be “signal coordinators”. Instead, they might appear in the bow-tie configuration with the purpose of being recruited for one task or the other dependent on the need of the cell, thus serving as multifunctional “knot” proteins.

Going further, we searched specifically for concrete examples of bow-ties that did not fit the “signal coordinator” role. One such example is presented in **Figure 4C**: Eleven “knot” proteins were located between the protein complexes “Ribonucleoprotein complex export from nucleus” and “Resolution of sister chromatid cohesion”. Neither the complexes nor the “knot” proteins are related to signal transduction. More specifically, most of the “knot” proteins were members of the nuclear pore complex (NPC), five of them members of the sub-nucleoporin complex Nup107-160 [22] and two of them were cyclins. NPC and Nup107-160 have been shown to participate in two different cellular processes: During cellular interphase, NPC and Nup107-160 form nuclear pores through which RNA and proteins are transported [23]. In contrast, during prometaphase, which is a mitotic phase where the nuclear envelope is dissolved, NPC and Nup107-160 participate in bipolar spindle assembly, wherein “Resolution of sister chromatid cohesion” is a required step [24]. Similarly, the cyclins CCNB2 and CDK1 forms a complex (<u>CPX-2007</u>), which has key roles in both “Resolution of sister chromatid cohesion” (<u>R-HSA-2500257</u>) and “Nuclear pore complex disassembly” (<u>R-HSA-3301857</u>). Thus, we argue that the NPC-related proteins and cyclins do not act as “signal coordinators” in this example. Instead, we argue that the bow-tie motif here is related to the multifunctionality of the “knot” proteins. Specifically, the bow-tie motif allows the “knot” proteins to be recruited by either of the two protein complexes when required.

This example of multifunctional “knot” proteins was not an isolated case. We also found protein UCH37 in a bow-tie motif between the proteasome complex and the human INO80 helicase complex (**Supplementary Figure 5A**). When bound to the proteasome, UCH37 assists in the proteolysis process by deubiquitylating proteins. In contrast, when bound to the human INO80 helicase complex, studies have proposed that UCH37 applies its deubiquitylation function on nucleosome remodeling and DNA repair [25,26]. Hence, the appearance of UCH37 in the protein interactome as a bow-tie motif seems directly linked to its multifunctionality. A third example was found in the members of the endonuclease complex, ERCC1-ERCC4-MUS81-SLX4 [27], together with members of “Replication protein A complex”, RPA1-RPA3 [28], suspended via bow-ties between the DNA repair complexes “Interstrand Crosslink Repair” and “HDR through Homologous Recombination” (**Supplementary Figure 5B**). The endonuclease complex plays a key role in forming and resolving Holliday junctions in these two processes [27]. Similarly, the “Replication protein A complex” is known to bind single-stranded DNA to prevent reannealing and it is therefore associated with many types of DNA repair [28,29]. Again, the multifunctional nature of these proteins seems to explain their position as “knots” in bow-tie motifs.

To assess whether the 294 “knot” proteins, as a group, were significantly enriched for multifunctional proteins, we ran a permutation test comparing the median number of associated GO biological processes (GO:BP) of 100,000 random sets of 294 proteins in the cDC1 interactome with the median number of associated GO:BP terms of the actual set of “knot” proteins. We used random protein sets with a degree distribution similar to that of the “knot” proteins to nullify the recognized positive relationship between a protein’s degree and its number of related GO:BP terms. We found that the “knot” proteins were associated with significantly more GO:BP terms, i.e. cellular functions, compared to random sets of proteins (**Figure 5,** *P* value = 1.5e-4). This finding, together with the previous examples of bow-tie motifs presented above, supports the hypothesis that some bow-tie motifs, which are not acting as “signal coordinators”, may assist or enable specific “knot” proteins in carrying out their multifunctional capabilities.

**Figure 5:**
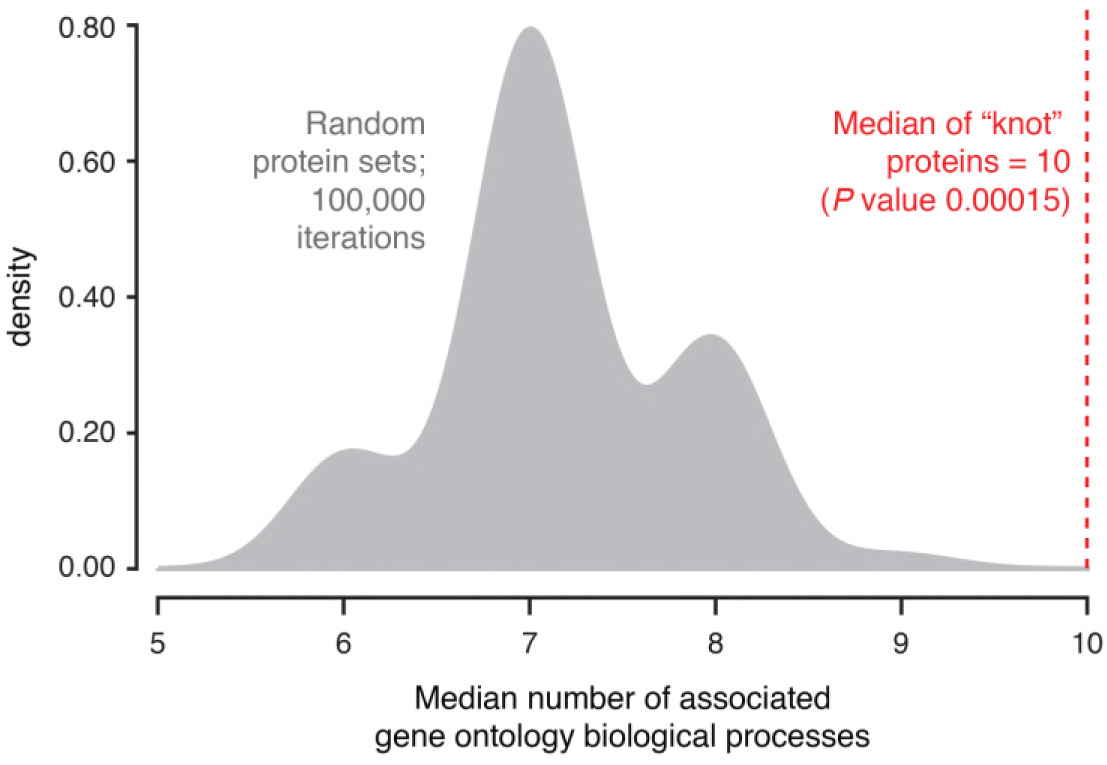
Significant association between bow-tie “knot” proteins and multifunctionality: Permutation test comparing the median number of associated GO:BP terms of the 294 identified “knot” proteins to 100,000 random protein sets of same size with similar node degree distribution. We found the “knot” proteins associated to a significantly higher number of GO:BP terms than expected by random (Median = 10, P value = 1.5e-4).

## Discussion

Cell type-specific protein interactomes are incredibly complex, often containing more than 10,000 proteins [30] that form dense interaction networks, typically with an average shortest path of 2-3 steps. This complexity makes the study of their topology very hard. Finding approaches to decode the topology and uncover topological trends, will facilitate actual use of the interactomes, for example for mapping perturbation propagation patterns or modeling context-dependent interactome dynamics.

We present a generic visualization and decoding approach that allows a detailed determination of the entire topology of very large networks. In this study, we applied it to the cDC1-specific protein interactome (n=8,568) to characterize its intermediate topological level, where protein complexes, and the topology that connects them, are found. We show that the bow-tie motif is a significant connector of cellular processes, especially fundamental processes common to many cell types. The bow-tie motif has previously been functionally characterized as a “signal coordinator” [16]. In this study, we add to the bow-tie motif’s repertoire of functional properties by proposing an alternative role, where the bow-tie topology allows the sharing of its “knot” protein between multiple functions, thus enabling multifunctionality.

We provide a novel approach to decode the topology of very large networks, using topological overlap matrices. Though the topological overlap has been applied before to illustrate hierarchy in smaller networks (n=195) [13,31], we further refine its use by applying the *weighted* topological overlap (wTO), developed by Zhang and Hovarth [18], to a network 45 times larger (n=8,568) and we present nearly 2,000 times more pairwise wTO calculations. We then demonstrate a novel and systematic way of reading the topology of a very large network, which is encoded in the wTO matrix, by linking known topological motifs to recognizable features in the matrix – and conversely. The topology of large networks, which could not be decoded otherwise due to their size, can hereby be examined in detail. Especially the identification of very large topological organizations of proteins (n∼100) is a unique feature of our approach.

The bow-tie is a well-known network topological motif [15–17]. In cell biology, however, it has been identified only in signaling-and metabolic networks [16,20,32,33]. Here, we greatly expand upon the bow-tie motif’s application in protein interactomes, showing that it is far more abundant than previously demonstrated, and that it populates functionally diverse areas of the protein interactome, many of which are not signaling networks. By presenting the bow-tie motif in the context of the intermediate topological level, we also show that the bow-tie motif not only connects upper and lower parts of signaling cascades, which is well-known [20,32], but that its main application may in fact be connecting distinct cellular processes. We find that the design of the bow-tie motif seems to allow “knot” proteins to be recruited by alternating cellular processes via their interaction fans when needed. The recruitment could be context dependent, changing between cellular states (**Figure 4C**). These results significantly reframe the bow-tie motif, revealing that it is more than a component of signaling networks. Such a setup may also aid in restricting genome size in terms of the number of proteins needed to drive cellular biochemistry. The human genome contains surprisingly few protein-coding genes, and the accepted number is decreasing in reference genome revisions, as some predicted genes and intron-free open reading frames are found to be non-coding [34]. The bow-tie motif may help keep the genome size down by enabling effective protein sharing.

Connecting cellular processes using the bow-tie motifs revealed a scaffold within the otherwise enigmatic cDC1 interactome, which was distinctly hierarchical. Hierarchy, as a network property, has previously been described in protein interactomes and large metabolic networks [12,13]. In those studies, the hierarchy was elegantly detected by plotting the clustering coefficient for each degree in the investigated networks. However, that approach did not reveal the actual topological motifs that generated the hierarchy and a knowledge gap thus remained. Given our observations, we propose that the hierarchical scaffold formed by the bow-tie motifs could, in part, fill this gap, as it evidently generates a hierarchy within the protein interactome.

## Methods

### Mice

C57BL6 mice were purchased from Janvier and maintained at the Biomedical Center (BMC), Lund University. Animal experiments were performed in accordance with the Lund/Malmö Animal Ethics Committee.

### Cell preparation

Cell suspensions of murine MLN were prepared as previously described [35]. Briefly, MLN were cut into small pieces and enzymatically digested with Collagenase IV (0.5 mg/ml, Sigma-Aldrich) and DNase I (12.5 mg/ml, Sigma Aldrich) for 45 min at 37°C while shaking and filtered prior to analysis. To pre-enrich the samples prior to cell sorting, B and T cells were labeled with biotinylated antibodies to CD19 and CD3, respectively, and depleted by using EasySep Streptavidin RapidSpheres (STEMCELL Technologies) according to manufacturer’s instructions.

### Flow cytometry analysis and cell sorting

MLN cell suspensions were stained with a cocktail of antibodies to CD45 (30-F11), CD3 (17A2), CD19 (6D5), B220 (RA3-6B2), TER-119 (TER-119), NK-1.1 (PK136), MHC II (IA/I-E) (M5/114.15.2), CD11b (M1/70), CD11c (N418), CD103 (M290), CD64 (X54-5/7.1), F4/80 (BM8), SIRPα (P84), XCR1 (ZET). CD11c^+^MHCII^hi^XCR-1^+^CD103^+^ cDC1 were flow cytometry cell sorted on a FACSAriaII (BD Biosciences) after exclusion of dead cells stained by fixable live/dead near-infrared dye (Life technologies) and cell aggregates (identified on FSC-A versus FSC-W scatterplots).

### Library preparation

Total RNA was isolated using the RNeasy Micro kit (QIAGEN). Briefly, frozen pellets of sorted cells were resuspended in RLT buffer. Buffer and cells were mixed for 30 sec through pipetting and left on the bench for 5 min to assure proper lysis. Extraction was performed according to manufacturer’s protocol with an on-column DNAse digestion step added after the first wash buffer step. RNA quality and quantity were measured using the 2100 BioAnalyzer equipped with RNA6000 Pico chip (Agilent Technologies). RNA was subjected to whole transcriptome amplification using Ovation RNA-Seq System V2 (NuGEN) kit, following the manufacturer’s protocol. This kit employs a single primer isothermal amplification (SPIA) method to amplify RNA into double stranded cDNA. Amplified cDNA samples were purified using the MinElute Reaction Cleanup kit (Qiagen) according to manufacturer’s instructions. Quantity and quality of the cDNA samples were measured using the 2100 BioAnalyzer equipped with DNA1000 chip (Agilent technologies) and the Nanodrop (ThermoFisher Scientific). Libraries to sequence with Illumina platform were constructed the Ovation Ultralow system V2 kit (NuGEN), following manufacturer’s instructions. 100ng of amplified cDNA were fragmented by sonication using a Bioruptor Pico (Diagenode). The sheared cDNA was end-repaired to generate blunt ends, then ligated to Illumina adaptors with indexing tags, followed by AMPure XP beads purification. The final size distribution of the libraries was evaluated using 2100 Bioanalyzer equipped with DNA1000 chip (Agilent technologies) and quantified using KAPA library Quantification Kit Illumina platforms (Kapa Biosystems).

### mRNA sequencing using Illumina

Equimolar amounts of each sample library (n=9) were pooled and the pools were used for cluster generation on the cBot with the Illumina TruSeq SR Cluster Kit (v3). Sequencing was performed on an Illumina HiSeq 1000 instrument using indexed 50 cycles single-read protocol and the TruSeq SBS v3 Reagents according to the Illumina HiSeq 1000 System User Guide. Image analysis and base calling resulted in BCL files, which were converted to FASTQ files with the CASAVA1.8.2 software. The resulting libraries had a length of 50 base pairs and 25-40 million reads per sample.

### RNA-seq data analysis

The quality of the FASTQ files was initially checked using FastQC [36]. The reads were aligned to the ENSEMBL mouse genome (GRCm38 release 89) using STAR version 2.4.2a [37] with default settings, which also quantified the gene expression. Gene expression files were preprocessed as recommended in Law *et al*. [38]. Briefly, genes with a log2 count per million (log2-cpm) of at least 1.0 in three out of nine samples were considered expressed. Raw counts were hereafter transformed using the *voom* function of the *limma* R package [39]. The pipeline produced a final table of 10,563 expressed genes across nine samples.

### Analysis of sorted immune cell RNA-seq data (public data)

FASTQ files associated with the public dataset E-GEOD-60424 [21] (ArrayExpress) were downloaded using the fastq-dump function from the sratoolkit (v. 2.8.2). High quality sequencing was confirmed using FastQC [36]. Alignment of the FASTQ files and gene expression quantification was performed as described above (**RNA-seq data analysis**). We only used the samples that were metadata tagged as “normal”, i.e. from healthy patients. Preprocessing of the files were also done as described above (**RNA-seq data analysis**).

### The cDC1-specific protein interactome

To construct the cDC1-specific protein interactome, we used the protein-protein interactome InWeb_IM [2] as foundation. InWeb_IM contains experimentally generated and scored physical PPIs from mainly *h*. *sapiens* (n=332,385), *m*. *musculus* (n=169,347) and *s*. *cerevisiae* (n=102,150) (PPI counts from [2]). Each PPI is benchmarked against a gold standard and scored accordingly, summed into a confidence score (cs). Only proteins expressed (defined above) in cDC1 cells were kept to generate the cDC1-specifc interactome. This method has previously been suggested in [5] and applied in [30]. Since InWeb_IM is indexed using UniProtKB/Swiss-Prot identifiers of *h*. *sapiens*, the 10,563 ENSMUSG identifiers of *m*. *musculus*, which represented the expressed genes in cDC1, were first mapped to *h*. *sapiens* ENSG identifiers (n=9,183) using ortholog pairing and then further mapped to *h*. *sapiens* UniProt/Swiss-Prot identifiers (n=8,984). The mapping was done using the R package *gProfileR* [40]. The final cDC1-specific interactome contained 8,568 expressed proteins, which had at least one PPI in the InWeb_IM database.

### The cDC1-specific interactome topology

To describe the topology of the cDC1-specific interactome, we calculated the pairwise wTO between all protein pairs. The wTO has previously been used in an unweighted form in Ravasz *et al*. [13] and been refined to its weighted form in Zhang *et al*. [18]. Given a protein *x* and a protein *y* in the interactome and with *a* symbolizing a connection strength between two proteins (0 ≤ *a* ≤ 1), the wTO was calculated using:

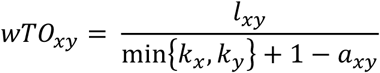

where 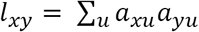 is the product of the connection strengths between *x* with *u* and *y* with *u*, where *u* is a common protein neighbor of both *x* and *y*, summed over all common protein neighbors of *x* and *y*. By neighbors, we refer to first order interactors of x and y only. The term 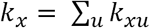 is the summed connection strength of protein *x* to all of its neighbors. For each protein pair *x* and *y*, this will produce a wTO between 0 and 1. Since we based our analysis on InWeb_IM, we used InWeb’s internal confidence score (0 ≤ *cs* ≤ 1) as a measure of inter-protein connection strength. The all proteins versus all proteins wTO matrix was constructed, resulting in an 8,568 x 8,568 symmetric wTO matrix (36.7 million unique pairwise comparisons).

### Matrix and network visualization

We produced a visualization of the cDC1 interactome topology in R by clustering the cDC1-specfic wTO matrix via *FlashClust* using the unweighted pair group method with arithmetic mean (UPGMA) [41], which resulted in a meaningful protein order, as previously demonstrated by Ravasz *et al*. [13]. We visualized the resulting matrix using the basic R function *image* with raster activated, creating a 73.4-megapixel image. All graph visualizations were done in Cytoscape [42].

### Test for multifunctionality using a permutation test

The 8,568 proteins in the cDC1-specific interactome were divided into four groups based on their degree quantile, i.e. their number of first-order interactions. Random protein sets (n=294) were then sampled 100,000 times from the cDC1-specific interactome (n=8,568), excluding the 294 “knot” proteins. We picked random proteins from degree quantile groups that corresponded to each of the 294 “knot” proteins. We hereby obtained random protein sets with a degree distribution similar to that of the “knot” proteins. For each random protein set, we then calculated the median number of GO:BP terms with which the proteins were associated. The generated *P* value was subsequently calculated by counting the number of random protein sets with a median GO:BP term count equal to or greater than that of the “knot” protein set (median = 10), divided by 100,000.

## Acknowledgements

RNA sequencing sample libraries were prepared by Julien Vandamme, PhD, Mucosal Immunology, Technical University of Denmark. RNA sequencing was performed at the Genomics Core Facility, KFB: Center of Excellence for Fluorescent Bioanalytics, University of Regensburg, Regensburg, Germany. Associate Professor Jose MG Izarzugaza, DTU Bioinformatics, Technical University of Denmark, provided expertise for the permutation test.

## Research funding

This work was supported by the Lundbeck Foundation (grant agreement 110145, RIMMI) and the Novo Nordisk Foundation (grant agreement NNF14CC0001 and grant agreement NNF17OC0027594, Challenge). Cristina Gomez-Casado was supported by a postdoctoral fellowship of Carl Tryggers Foundation.

## Supplementary Material

### Supplementary Figure 1

**Supplementary Figure 1:**
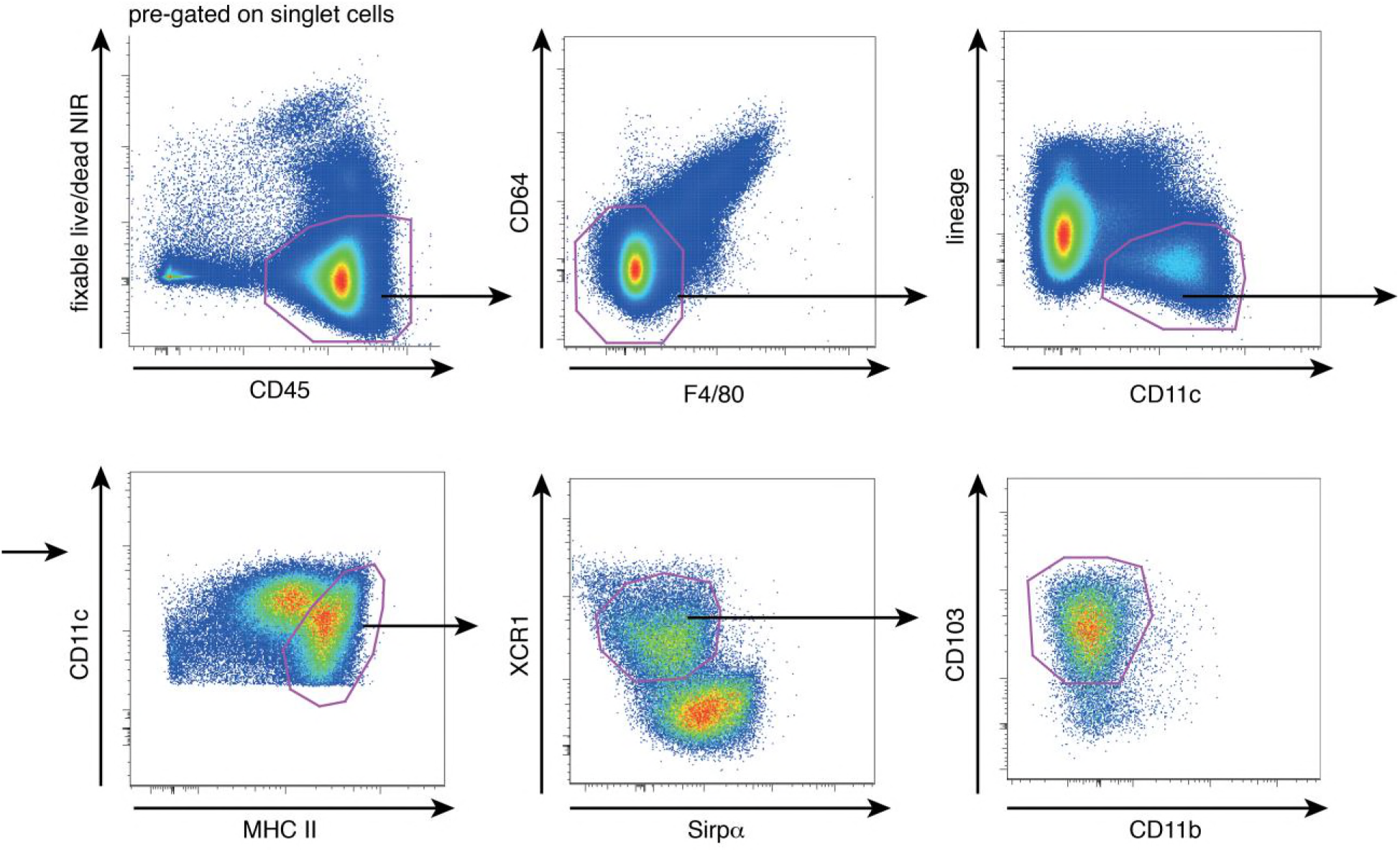
Flow cytometry gating strategy used to identify migratory CD103+ cDC1 cells from murine mesenteric lymph nodes. Lineage; CD3, CD19, B220, NK −1.1 and TER-119.

### Supplementary Figure 2

**Supplementary figure 2:**
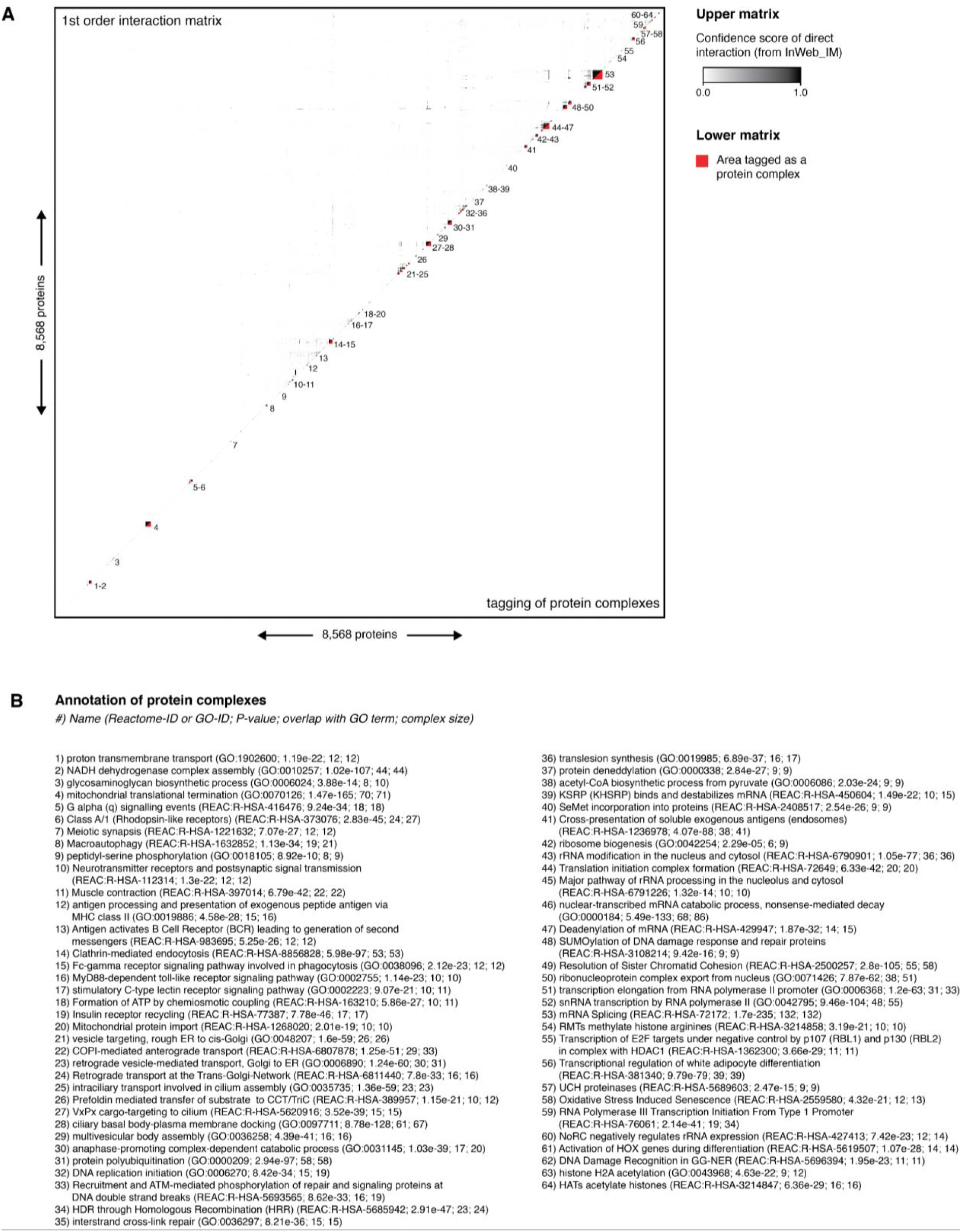
Tagging of protein complexes. (A) The 8, 568 × 8, 568 matrix showing the confidence score (0.0-1.0), as given by InWeb_IM, of all direct PPIs in the cDC1 interactome similarly to the upper part of Figure 2A. We constructed a script that traversed the diagonal of this c DC1 matrix (upper matrix) and tagged all protein complexes which we marked red and numbered (lower matrix). (B) All protein complexes annotated with a GO:BP term or Reactome pathway name, depending on which of those that were most significant.

### Supplementary Figure 3

**Supplementary figure 3:**
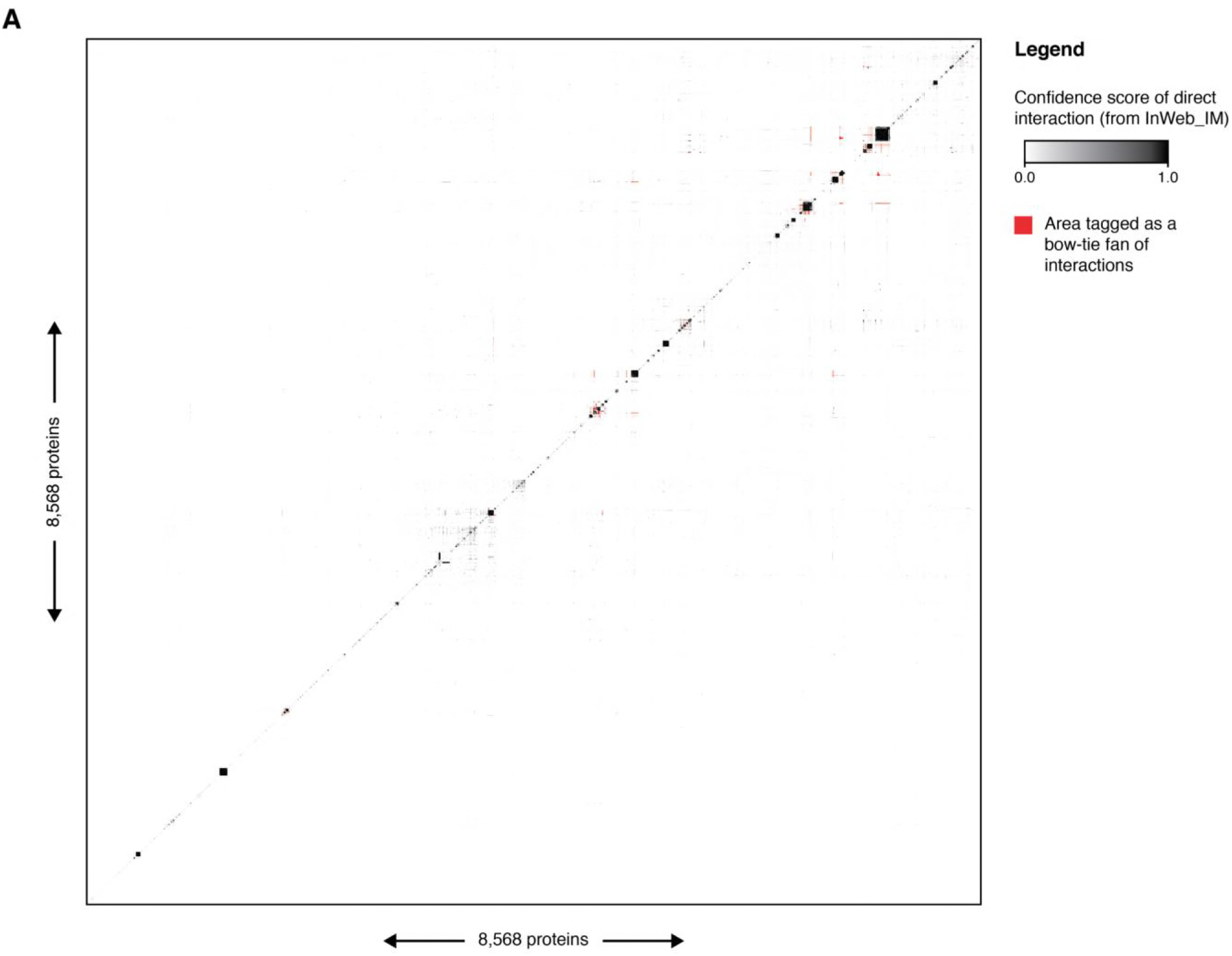
Tagging of bow-tie interaction fans in the cDC1 interactome. The 8, 568 × 8, 568 matrix showing the confidence score (0.0-1.0), as given by InWeb_IM, of all direct PPIs in the c DC1 interactome similarly to the upper part of Figure 2A. We constructed a script that tagged all bow-tie interaction fans defined as (i) located between two protein complexes and (ii) having 9/10 consecutive interactions on a vertical or horizontal line in the matrix with a confidence score of 1.0, thus, allowing 1/ 10 of the PPIs to be below 1. 0. These markings appear as “ dashes” in the matrix and are colored red.

**Supplementary Table 1:**
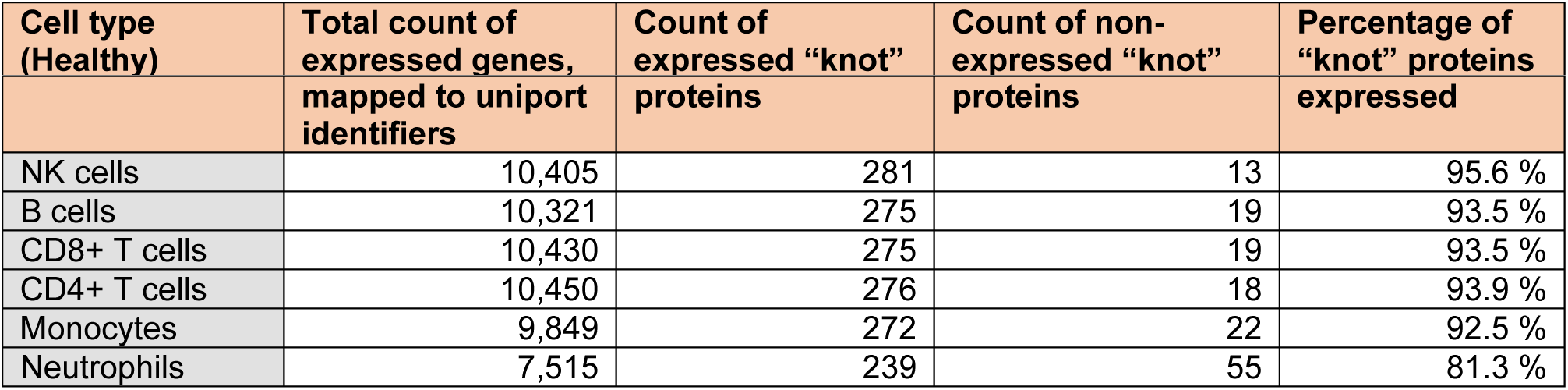
The majority of cDC1 “knot” proteins were expressed in six other human immune cells from blood.

### Supplementary Figure 4

**Supplementary figure 4:**
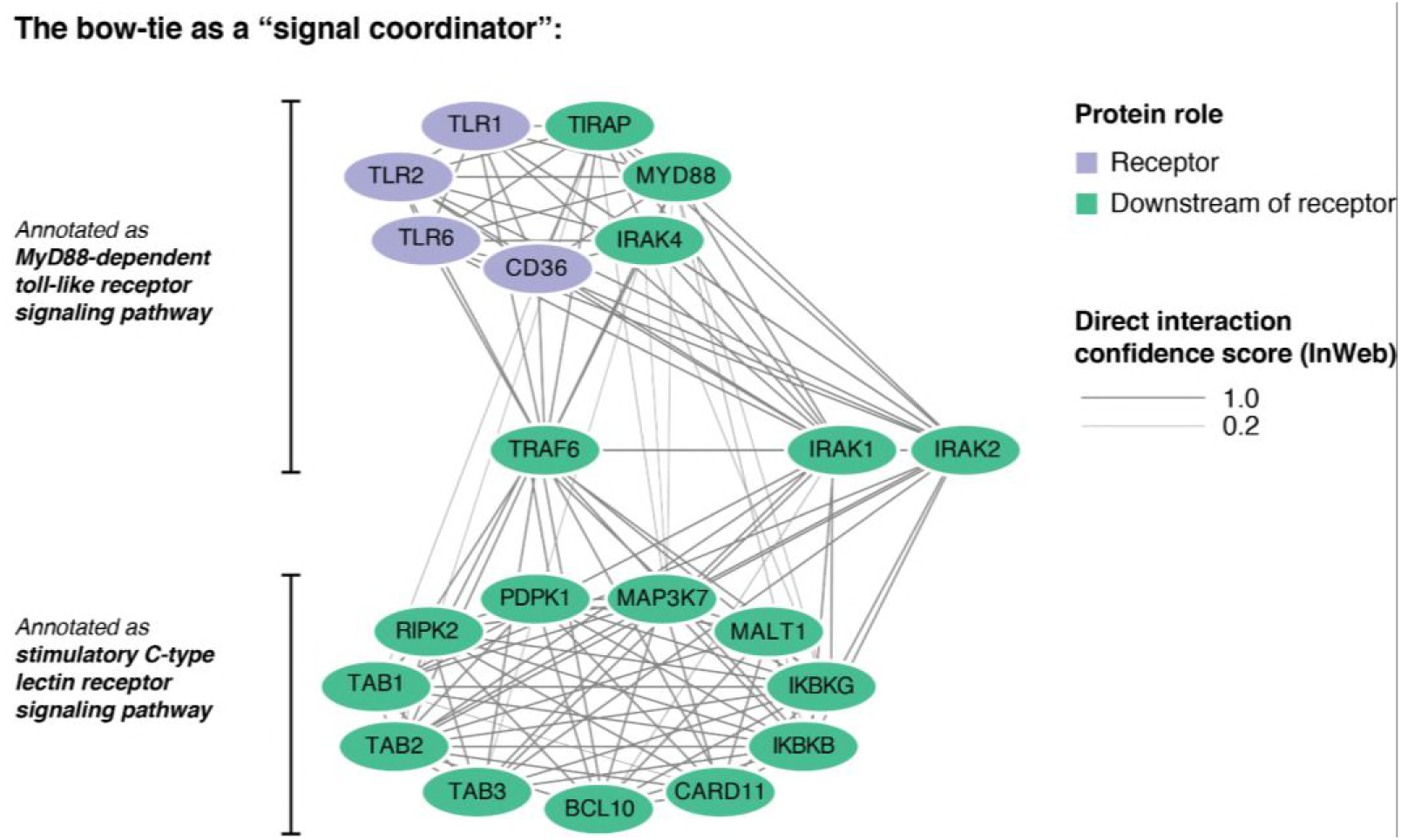
The functional role of bow-tie motifs as “ signal coordinators”. Network consisting of toll-like receptor (TLR) signaling proteins containing a bow-tie, TRAF6, and two potential bow-ties, IRAK1 and IRAK2 (although we could not definitively confirm them as “ bow-ties”). TRAF6 and IRAK1 have previously been described as bow-tie “ knot” proteins [20]. TRAF6, a signal transducer factor, carries signals from the TLRs (upper complex) to downstream pathways (lower complex), illustrating an example of the “ s ignal coordinator” role.

### Supplementary Figure 5

**Supplementary figure 5:**
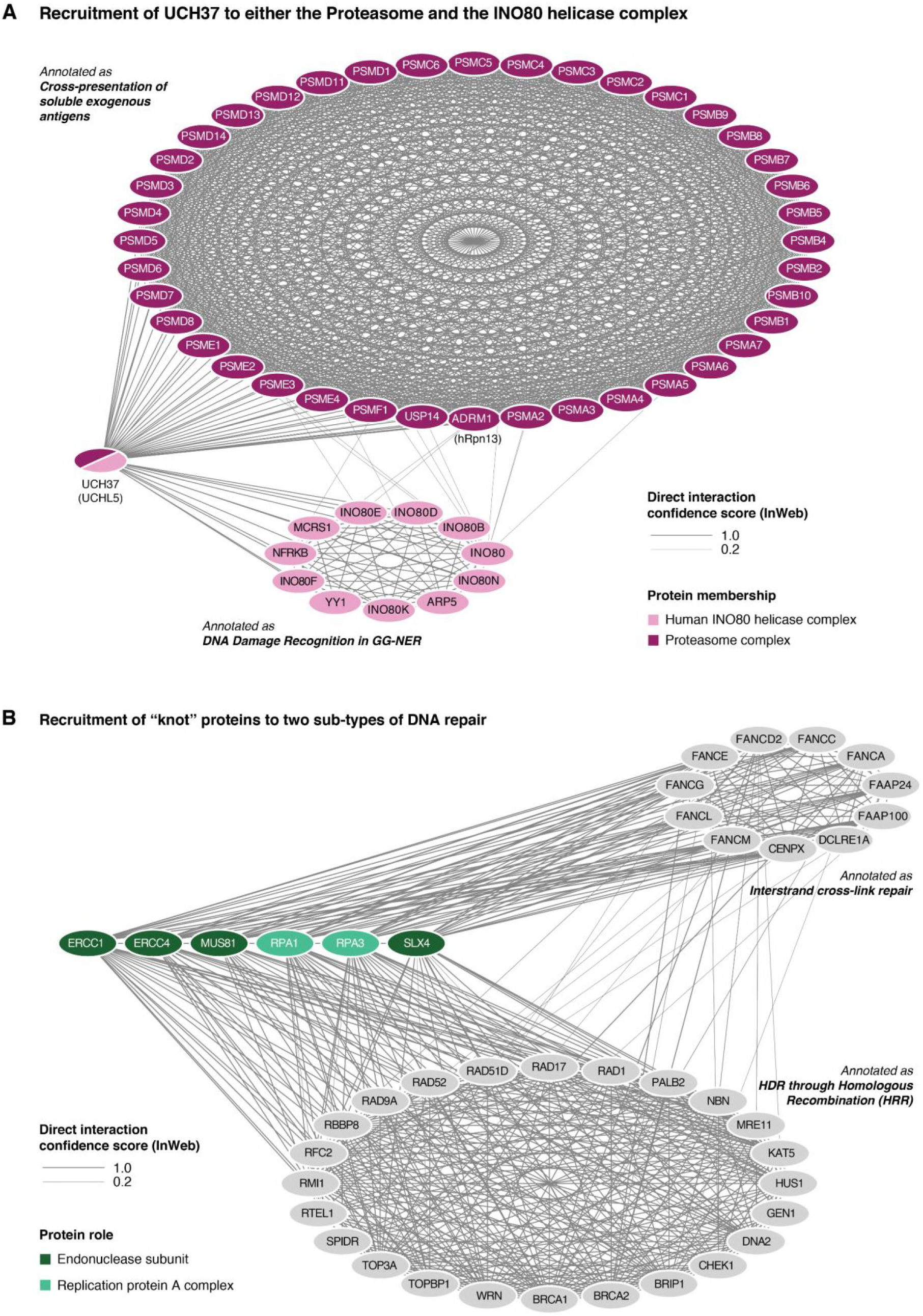
Bow-tie motifs enable the multifunctionality of “ knot” proteins: (A) The enzyme UCH37 applies its molecular function of deubiquitylation in two different setting s: as a member of the proteasome complex and as an associate of the INO80 complex. Suspended in a bow-tie motif between the two protein complexes, UCH37 can alternate between the two cellular processes when required. **(B)** Similarly, the endonuclease proteins (ERCC1, ERcc4, MUS81 and SLX4) and replication protein complex A (RPA1 and RPA3) are known to take part in both “Interstrand crosslink repair” and “ HDR through homologous recombination”. The bow-tie motifs appear to be an integral part of these proteins’ multifunctional capabilities, allowing them to switch between the two types of DNA repair.

